# Rapid optogenetic clustering of a cytoplasmic BcLOV4 variant

**DOI:** 10.1101/2023.09.14.557726

**Authors:** Zikang Huang, William Benman, Liang Dong, Lukasz J. Bugaj

## Abstract

Protein clustering is a powerful form of optogenetic control, yet there is currently only one protein —Cry2—whose light-induced clustering has been harnessed for these purposes. Recently, the photoreceptor BcLOV4 was found to form protein clusters in mammalian cells in response to blue light, although clustering coincided with its translocation to the plasma membrane, potentially constraining its application as an optogenetic clustering module. Herein we identify key amino acids that couple clustering to membrane binding, allowing us to engineer a variant of BcLOV4 that clusters in the cytoplasm and does not associate with the membrane in response to blue light. This variant, BcLOVclust, clustered over many cycles with dramatically faster clustering and de-clustering kinetics compared to Cry2. The magnitude of BcLOVclust clustering could be strengthened by appending an intrinsically disordered region from the fused in sarcoma (FUS) protein, or by optimizing the fluorescent protein to which it was fused. BcLOVclust retained the temperature sensitivity of BcLOV4 such that light induced clustering was transient, and the rate of spontaneous declustering increased with temperature. At low temperatures, BcLOVclust and Cry2 could be multiplexed in the same cells, allowing light control of independent protein condensates. BcLOVclust could also be applied to control signaling proteins and stress granules in mammalian cells. Thus BcLOVclust provides an alternative to Cry2 for optogenetic clustering and a method for multiplexed clustering. While its usage is currently suited for organisms that can be cultured below ∼30 °C, a deeper understanding of BcLOVclust thermal response will further enable its use at physiological mammalian temperatures.

**Graphical abstract:** 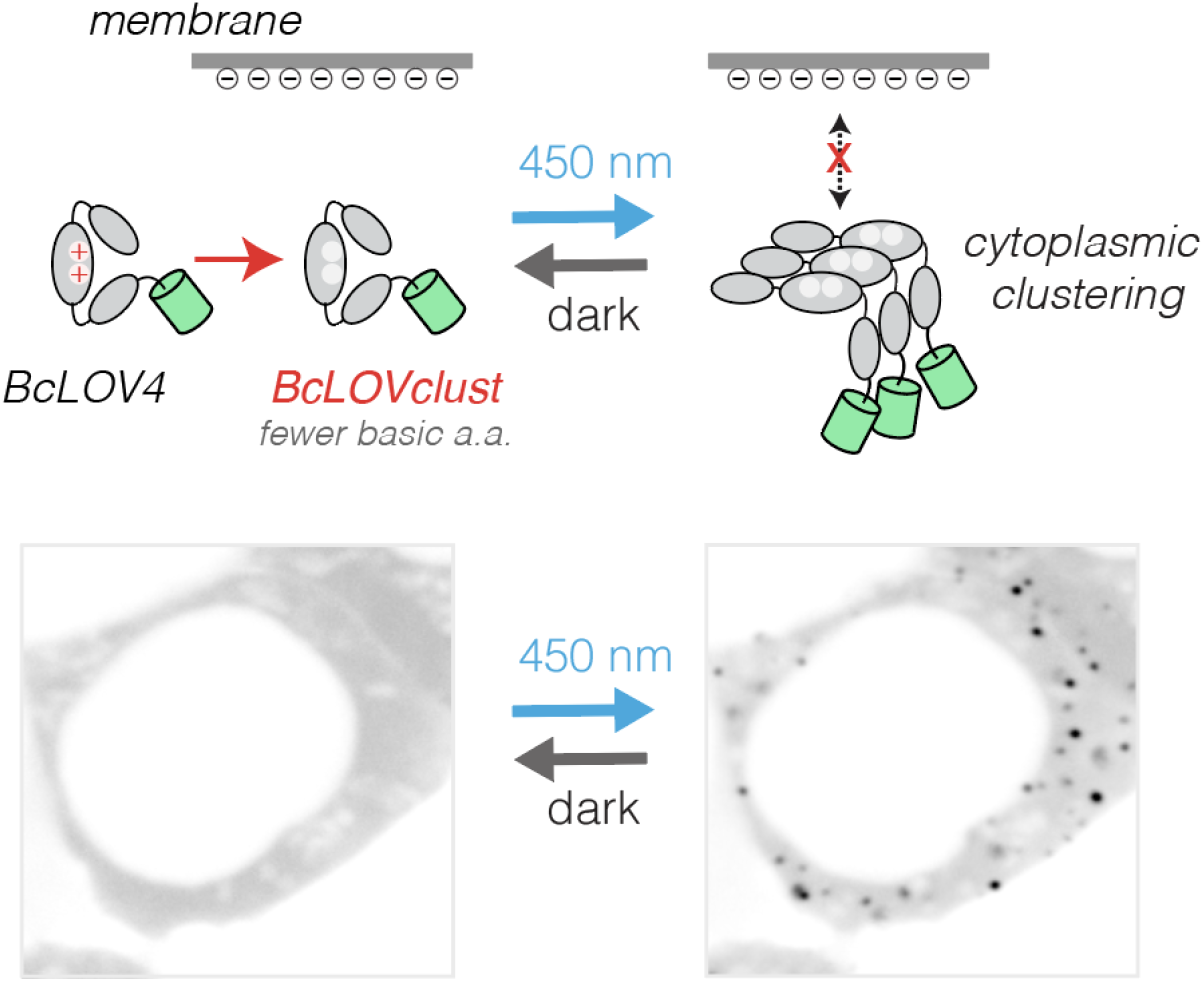

**Highlights:**

1. Light-responsive clustering of BcLOV4 can be decoupled from its membrane association
2. BcLOVclust clusters in the cytoplasm with faster ON and OFF kinetics than Cry2
3. BcLOVclust and Cry2 can be multiplexed in the same cell
4. BcLOVclust can control RhoA activity and stress granule formation

## Introduction

Protein clustering is crucial for a variety of cell behaviors across the domains of life [1]. Optogenetic protein clustering has provided tools for direct and dynamic study of such clusters with optical precision in living cells and animals. Such methods have allowed control of receptor tyrosine kinases in normal physiology and in cancer [2–5], stem cell development[6,7], calcium signaling and insulin secretion [8], biomolecular condensates [9,10], compartmentalization of biosynthetic pathways [11], and mammalian cell death [12], among many other applications.

Remarkably, despite the utility of optogenetic clustering, there remains primarily one protein whose native light-induced clustering has been leveraged for this purpose: *A. thaliana* Cryptochrome 2 (hereafter referred to as Cry2)[13]. To apply this one protein to a broad range of applications, Cry2 has been modified to tune its clustering properties. For example, the CRY2olig and Cry2clust variants provided stronger and faster light-induced clustering than wt Cry2 [14,15]. Separately, clustering could be enhanced with particular fluorescent proteins or the addition of intrinsically disordered regions (IDRs) [9,14].

Despite these advances, the lack of alternatives to Cry2 presents several challenges. The declustering kinetics of Cry2 are relatively slow, lasting ∼10s of minutes or greater depending on the fluorophore, and variants with enhanced clustering show slower kinetics [14]. Such kinetics prevent the use of Cry2 for certain applications, for example for studying the rapid dynamics of T cell receptor signaling [16–18]. Additionally, it is currently challenging to control multiple distinct clusters in the same cells, for example to better understand interactions and dynamics between multiple condensed species [19], [20]. Although non-Cry2-based optogenetic clustering methods could be applied, these require multiple components and stoichiometric tuning [21], complicating their use.

Recently, the BcLOV4 photoreceptor was found to form light-induced homo-oligomers at the plasma membrane, representing a possible alternative to Cry2 for optogenetic clustering [22,23]. Moreover, activation and inactivation kinetics were faster than for Cry2, with translocation/clustering occurring in seconds and dark reversion within a few minutes of dark adaptation [22]. However, BcLOV4 clustering happened concurrently with membrane translocation, potentially constraining its downstream applications. It is currently not clear how clustering and membrane binding are separately encoded in BcLOV4 sequence, nor whether cytoplasmic clustering can occur in the absence of its membrane binding.

In this work, we report an engineered variant of BcLVO4 called BcLOVclust that reversibly clusters in the cell cytoplasm without translocation to the membrane. BcLOVclust was generated by testing the hypothesis that membrane binding — but not clustering — required basic residues in the small amphipathic helix (AH1) downstream of the LOV domain. Similar to Cry2, BcLOVclust clusters across a wide range of expression levels, and clustering can be modulated by appending intrinsically disordered regions (IDRs) or by choice of fluorescent protein [9,14,24]. BcLOVclust retained the fast on- and off-kinetics as well as a temperature-sensitivity previously described for wtBcLOV4, and could be multiplexed with Cry2 to control independent clusters in the same cell. Finally, BcLOVclust could be used in the context of protein fusions to control RhoA activity and stress granule formation in mammalian cells.

## Results

### Isolation of clustering from membrane binding by removal of positive charges

BcLOV4 both translocates to the plasma membrane and forms membrane-associated clusters within seconds of blue light stimulation [22,23] (**Figure 1A**). Because membrane binding can be mediated through electrostatic interactions between positively charged amino acids and negatively charged phospholipids, we asked whether mutation of specific basic amino acids could eliminate membrane binding but retain clustering [22,25] (**Figure 1B**). We identified six lysines and one arginine in an extended amphipathic helix region (AH1e) as candidate amino acids that might contribute to membrane association, and we generated mutants with progressively more of these residues mutated to alanines (**Figure 1C**). When expressed in HEK 293T cells, all mutants were diffused in the cytoplasm in the dark and remained responsive to blue light, though with notable differences in localization (**Figure 1D-F, Supplementary Movie 1**). Mutants with fewer positive charges showed progressively less membrane localization yet retained their ability to cluster. Whereas wt BcLOV4 translocates and clusters almost instantaneously, the 2K->A mutant showed initial clusters both at the membrane and cytoplasm, with most cytoplasmic clusters eventually localizing to the membrane. The 4K->A mutant showed dramatically less membrane binding than 2K->A, though clusters were still visibly associated with the cell periphery. Finally, the 6K1R->A mutant showed no membrane binding (**Figure 1E, Supplementary Movie 1**). Clustering of the 6K1R mutant was somewhat diminished relative to the other mutants, suggesting that the basic residues tested may also partially contribute to clustering potential **(Figure 1F)**. Hereafter, we refer to the 6K1R->A mutant as BcLOVclust, a variant of BcLOV4 that clusters in the cytoplasm in response to blue light.

**Fig 1.**
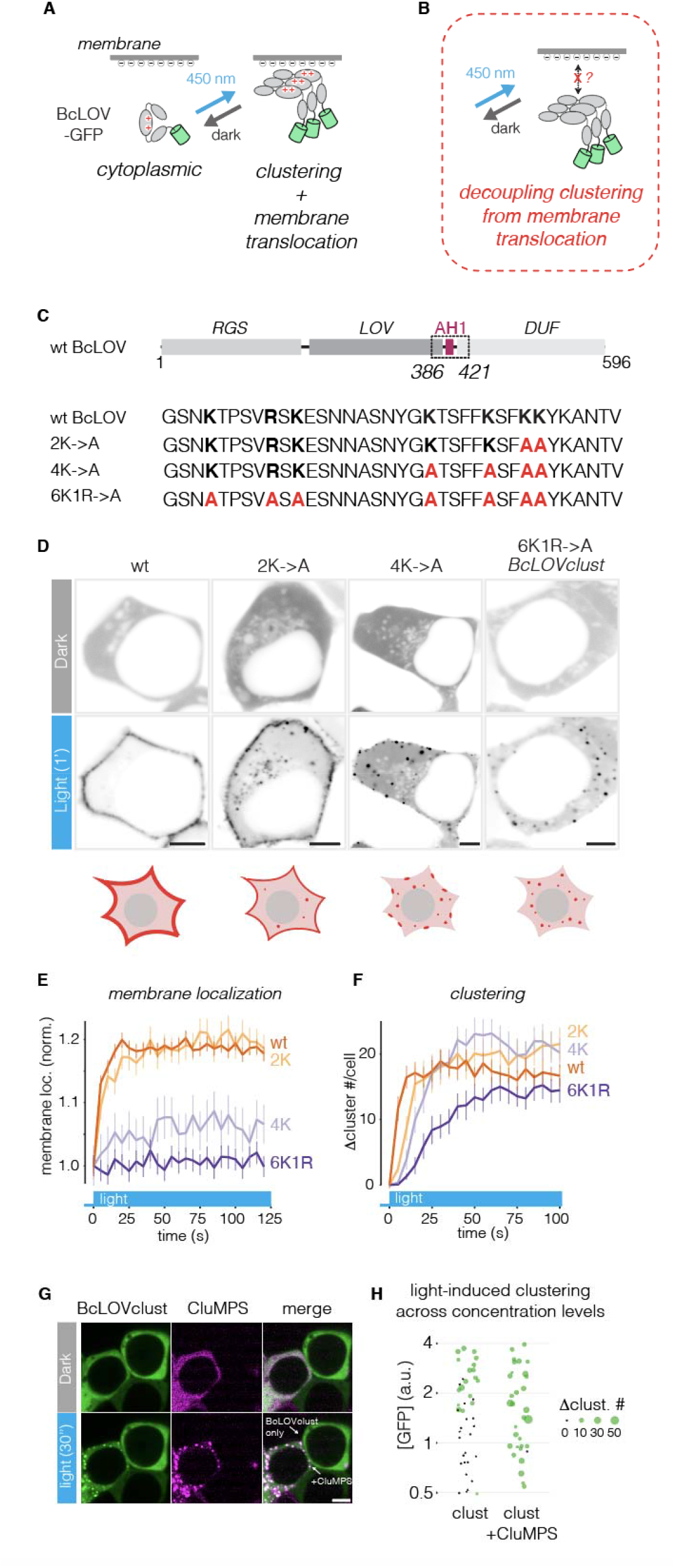
Isolation of BcLOV4 clustering from membrane binding by removal of positive charges around amphipathic AH1 region. a. BcLOV4 is a photoreceptor that both clusters and translocates to the plasma membrane upon stimulation with blue light. Membrane association was hypothesized to occur at least in part through electrostatic interactions between the AH1 region and negatively-charged membrane phospholipids. b. We tested whether removal of the appropriate positive charges would remove membrane association but retain light-induced clustering. c. Top: domain representation of BcLOV4, with extended AH1 (AH1e) region outlined. Bottom: amino acid sequence of wt AH1e and three mutants with progressively more removal of positive charges. d. Representative images of the distribution of BcLOV4 and mutants before and 1 min after light stimulation in HEK 293T cells. Cells were stimulated with a 488 nm laser every 5 seconds at 29 °C. e. Quantification of light-induced membrane association of BcLOV4 and mutants from data shown in (d). Membrane fluorescence is normalized by whole-cell fluorescence, and further normalized to the first time point (dark state). Data show mean +/− SEM of >50 cells per construct. f. Quantification of light-induced clustering of BcLOV4 and variants from data shown in (d). Data show mean +/− SEM of 15 cells per construct. g. Representative images show two cells with equivalent expression of BcLOVclust, one of which is co-transfected with a GFP-targeting CluMPS reporter. Whereas clusters are too small to see at this expression level, CluMPS amplifies and reveals their presence. Cells were stimulated with a 488 nm laser every 5 seconds at 27 °C. h. Comparison of clustering as a function of BcLOVclust-GFP expression level, as shown in (g). Clusters cannot be visualized below a threshold level. However, co-transfection with CluMPS reveals clusters at even these low expression levels, suggesting that BcLOVclust forms small light-induced clusters in this expression regime. Each data point is a single cell. Scale bar, 10 µm.

BcLOVclust displayed clustering in a concentration-dependent manner, where clusters were apparent only in cells with high levels of expression, similar to observations with Cry2 (**Figure 1G**) [24]. However, clustering can occur even if large fluorescent aggregates are not observed, for example because small clusters do not concentrate sufficient amounts of fluorescent protein to rise above the fluorescence background [24]. To ask if clustering occurred in low-expressing cells, we co-expressed BcLOVclust-GFP with a GFP-targeting CluMPS reporter (LaG17-mCh-HOTag3), which amplifies small clusters of GFP to enhance their detection, and we quantified clustering after light stimulation. Indeed, CluMPS revealed clustering in cells across all expression levels tested, including at low levels where clusters were not observed without CluMPS (**Figure 1G,H**)

### BcLOVclust has fast clustering kinetics

We next characterized the kinetics of BcLOVclust clustering. We expressed BcLOVclust-GFP or Cry2-GFP and we measured cluster formation and dissociation after light stimulation or removal, respectively. In cells with comparable expression levels, BcLOVclust-GFP showed substantially more rapid cluster formation compared to Cry2-GFP (T_1/2 (BcLOVclust)_ = 27.3 s, T_1/2 (Cry2)_ = 42.8 s) (**Figure 2A, S1A**), though the number and size of Cry2 clusters surpasses those of BcLOVclust at steady-state(**Figure S1B-C**). BcLOVclust-Venus also showed faster declustering compared to Cry2-Venus, with a ∼8-fold lower half-life of clusters after light removal (T_1/2 (BcLOVclust)_ = 2.5 min, T_1/2 (Cry2)_ = 19.1 min) (**Figure 2B**). These faster kinetics suggested that BcLOVclust might outperform Cry2 in tracking rapid pulses of input stimulation. Indeed, when stimulated in patterns of 2 min ON and 10 min OFF, BcLOVclust-GFP faithfully tracked light stimulation, with effectively full cluster dissociation before the subsequent stimulation. By contrast, Cry2-GFP behaved like a low-pass filter, maintaining a roughly constant cluster number over time because clusters persisted from the previous stimulation (**Figure 2C-E**). Thus, BcLOVclust allows for cytoplasmic clustering with more rapid dynamics than achievable with Cry2.

**Fig 2.**
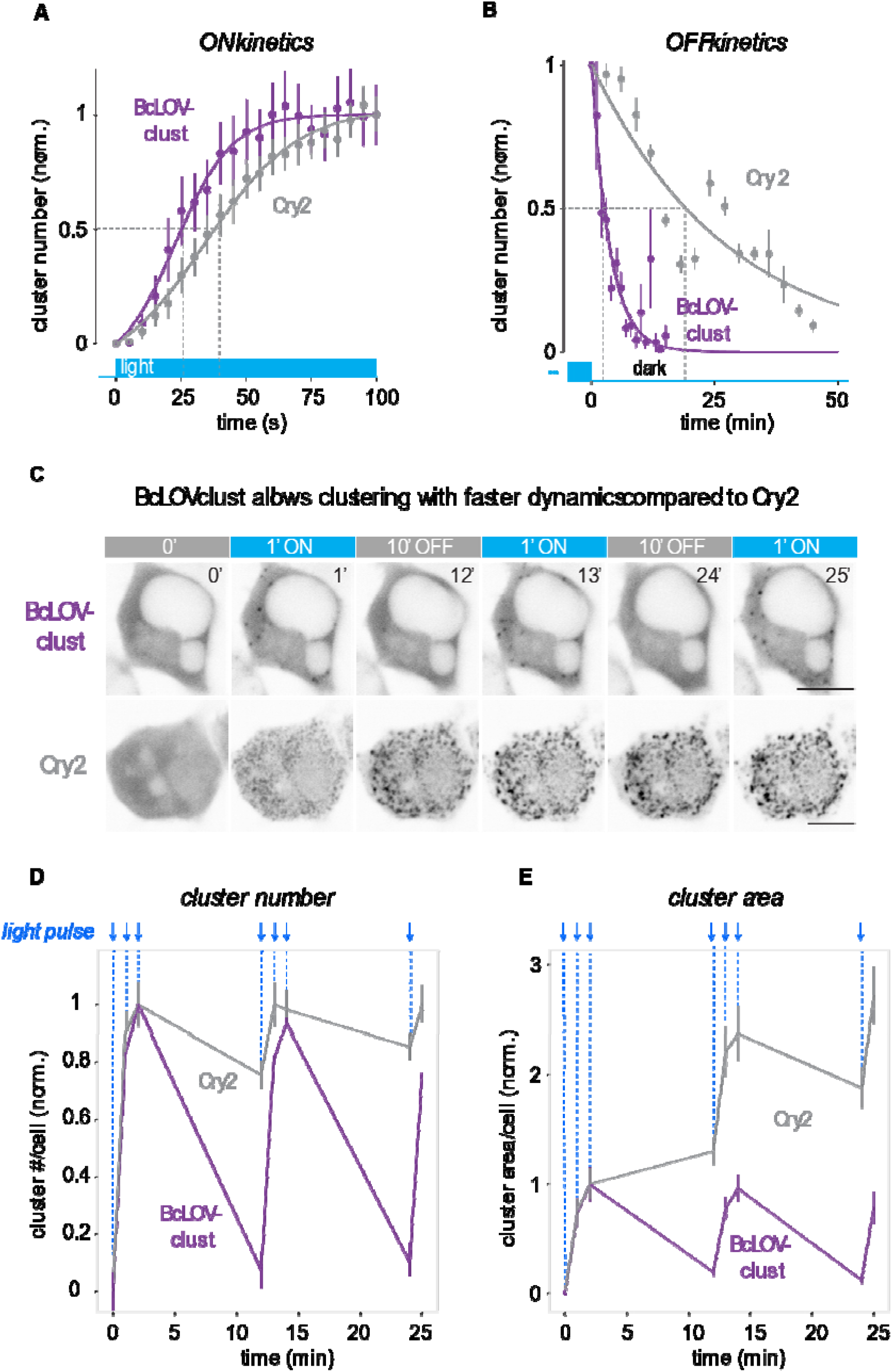
Characterizing fast ON and OFF kinetics of BcLOVclust. a. Quantification of clustering kinetics of BcLOVclust-GFP and Cry2-GFP upon blue light stimulation. Cells were exposed to 488 nm laser light every 5 seconds under a confocal microscope at 29 °C. Cluster number was normalized between min and max. Data show mean +/− SEM of more than 15 cells per construct. b. Quantification of declustering kinetics of BcLOVclust-mVenus and Cry2-mVenus. To avoid potential artifacts associated with optogenetic stimulation from mVenus imaging, cells in separate wells were imaged at the indicated time points after light stimulation (pulse of 488 nm light per minute for 3 minutes at 25 °C). Cluster number was normalized between min and max. Cells with equivalent fluorescence levels were measured. Data show mean +/− SEM of 5-10 cells per time point for each construct. See **Methods** for details on end-point pseudo-time-course analysis. c. Representative images showing pulsatile activation of BcLOVclust-GFP or Cry2-GFP. The two cells shown have similar expression levels. d. Quantification of cluster number (d) and area (e) of BcLOVclust-GFP and Cry2-GFP upon pulsatile light stimulation, with illumination from the 488 nm laser light source. Light pulses are indicated by blue arrows. For experiments depicted in (c-e), cells were stimulated and imaged at 25 °C. Data show mean +/− SEM of more than 15 cells per construct. Scale bar, 10 µm.

### Modulation of BcLOVclust with intrinsically disordered regions and fluorescent proteins

Despite its fast clustering kinetics, BcLOVclust forms fewer clusters than Cry2 at equal expression levels, potentially limiting its application. Inspired by prior studies with Cry2, we asked whether fusion of intrinsically disordered regions (IDRs) might promote stronger light-induced clustering [9]. Appending the low-complexity domain from the FUS protein (FUS-LC) to the C-terminus dramatically increased the number of light-induced clusters that appeared in each cell (**Figure 3A-C**). FUS-LC also altered the kinetics of clustering, with slightly faster clustering (T_1/2 (BcLOVclust)_ = 19.7 s, T_1/2 (BcLOVclust-FUS)_ = 14.1 s) and slightly slower declustering kinetics (T_1/2 (BcLOVclust)_ = 1.4 min, T_1/2 (BcLOVclust-FUS)_ = 3.0 min) (**Figure 3D-E**). Addition of FUS-LC also yielded more robust clustering under low expression levels (**Figure 3F**). We also tested an N-terminal fusion of FUS-LC but observed that, although clustering was similarly enhanced, the FUS-BcLOVclust clusters formed fiber-like aggregates as opposed to puncta at higher expression levels (**Figure S2**). We thus exclusively used C-terminal fusions of FUS-LC for the rest of this work. Additionally, we tested whether certain fluorescent proteins (FPs) could impact the extent of BcLOVclust clustering, as observed previously for Cry2 [14]. Whereas fusions to GFP or mVenus clustered more robustly, fusions to mCherry only had minimal response upon light stimulation (**Figure 2, S3**). Thus for the majority of the manuscript, we used GFP fusions unless otherwise specified.

**Fig 3.**
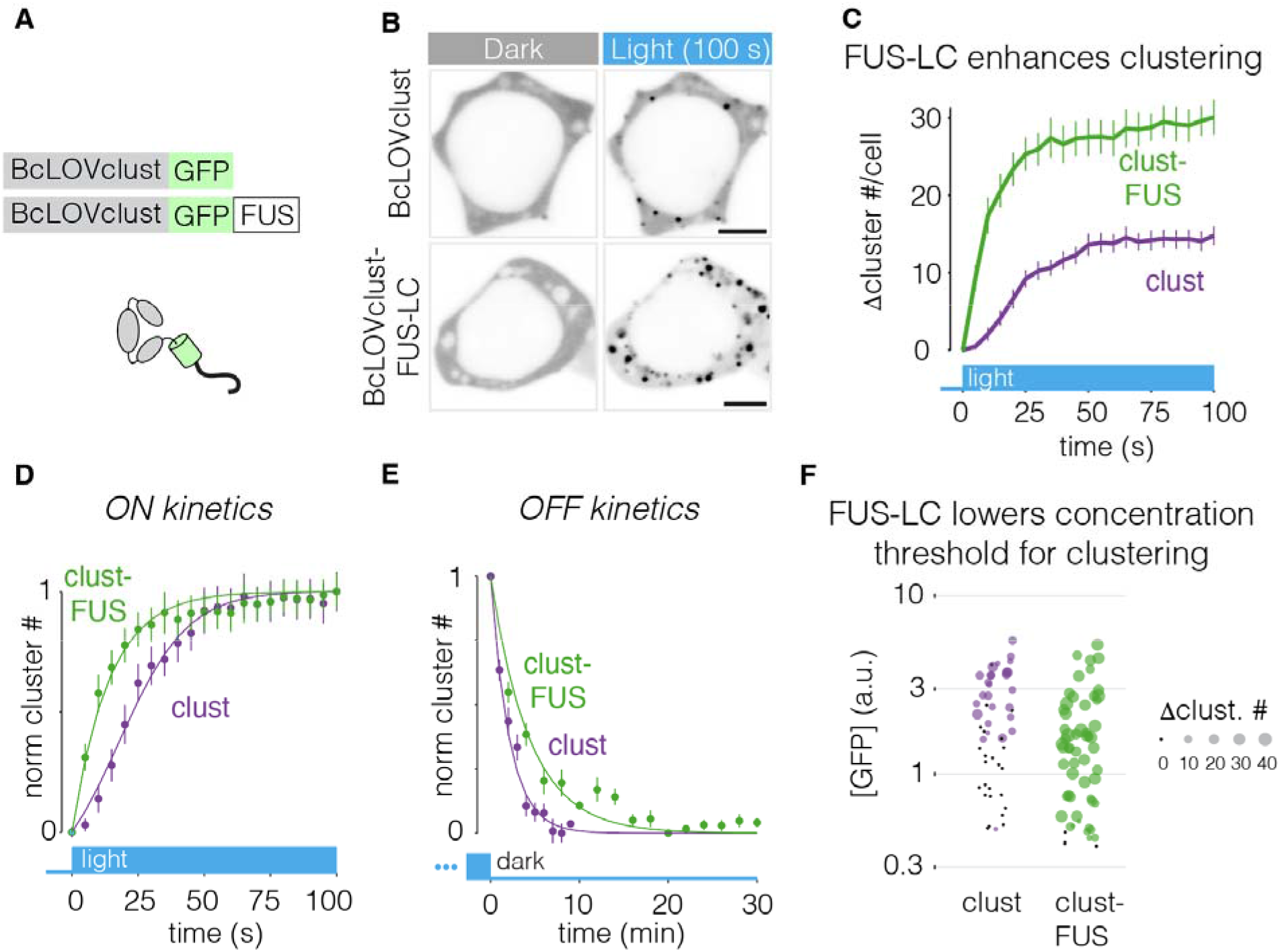
Magnitude of BcLOVclust clustering can be tuned by FUS-LC. a. Schematic of BcLOVclust-GFP fused to the FUS low complexity domain (FUS-LC). b. Representative images of light induced clustering of BcLOVclust-GFP or BcLOVclust-GFP-FUS. c. Quantification of clustering of constructs depicted in (a) at 25 °C. Data show mean +/− SEM of ∼15 cells per construct. d. FUS speeds the kinetics of BcLOVclust clustering. Plot shows data from (c) normalized between min and max. e. FUS moderately slows kinetics of declustering. Cluster number was normalized between min and max. Data show mean +/− SEM of 5-10 cells per time point for each construct. Data was obtained in the same manner as in Figure 2B. f. Comparison of visible clustering as a function of expression level. FUS-LC lowers the expression level at which BcLOVclust clusters can be observed. The increase in cluster number after 100s of stimulation is shown. All cells in this figure were stimulated every 5s at 27 °C. Scale bar, 10 µm.

### Temperature-dependence of BcLOVclust clustering

We previously found that optogenetic activation of BcLOV4 depended on temperature, such that light-dependent membrane translocation could be sustained below 30 °C, but translocation was only transient above 30 °C despite constant illumination [26]. Moreover, the rate of dissociation from the membrane increased as temperatures increased above 30 °C [26]. We thus asked whether clustering of BcLOVclust retained this temperature sensitivity. Indeed, clustering of BcLOVclust was strongest when cells were kept at lower temperatures (27 °C) and weakened as temperatures were raised to 30 °C (**Figure 4A-B**). No visible clustering was observed at or above 31 °C. While small clusters could still be detected at this temperature with CluMPS, these clusters could no longer be seen at 32°C (**Figure 4A-B, S4A-B**).

**Fig 4.**
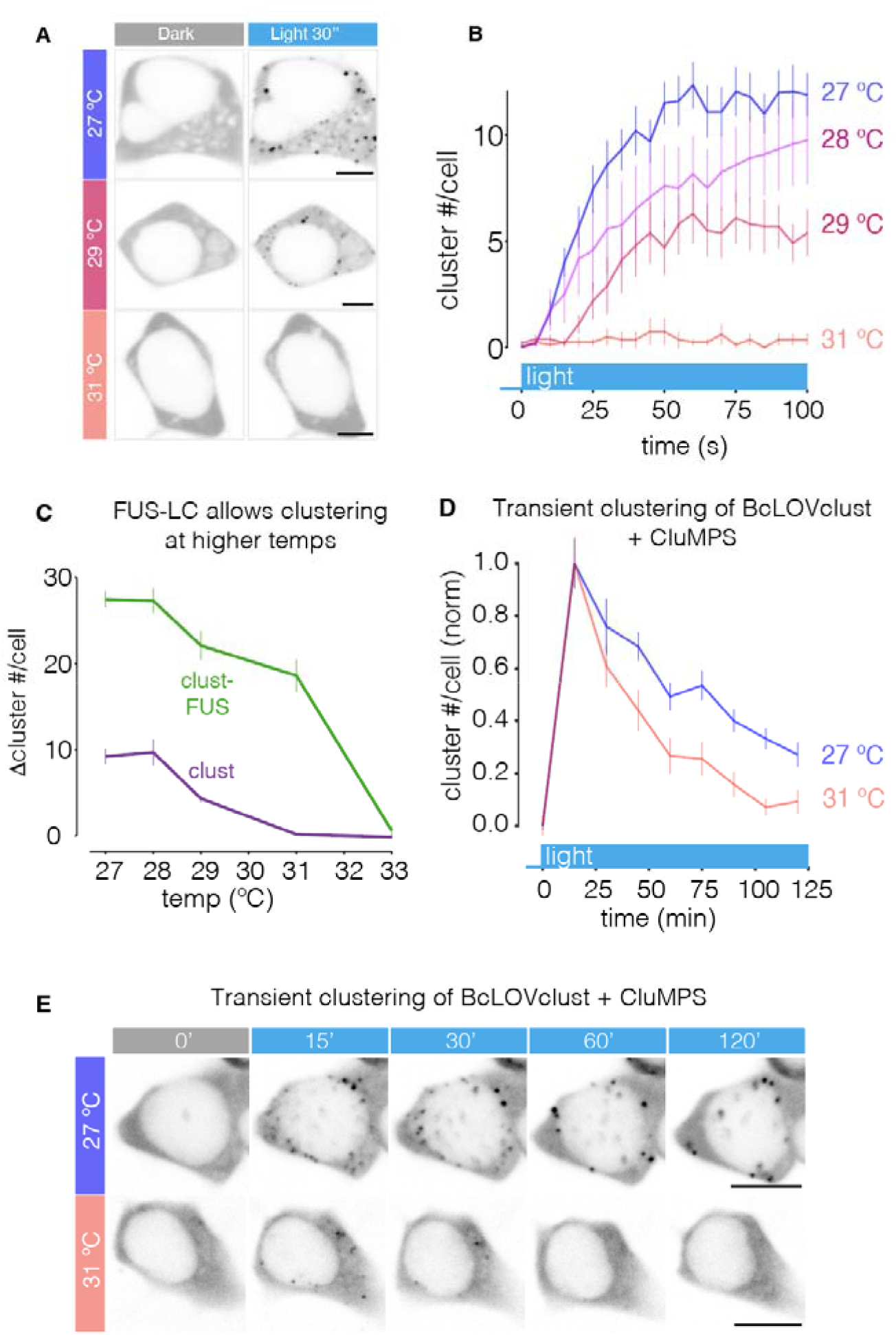
BcLOVclust clusters in a temperature-dependent manner. a. Representative images of BcLOVclust clustering under different temperatures. b. Quantification of clustering under the indicated temperatures. Data show mean +/− SEM of ∼10 cells in each condition. c. Comparison of steady-state cluster formation (100s) of BcLOVclust vs BcLOVclust-FUS at the indicated temperatures. Data show mean +/− SEM of ∼10 cells for each construct. d. BcLOVclust transiently clusters at elevated temperatures, as depicted in the presence of a CluMPS reporter to visualize small clusters. Cluster number of CluMPS channel is normalized to the 15 min time point. e. Representative images of date in Figure 4D showing transient clustering of BcLOVclust at the indicated temperatures. BcLOVclust-GFP channel is shown. Scale bar, 10 µm.

Our prior work showed that the FUS-LC domain could prolong light-induced membrane localization of BcLOV4 at 37 °C [23]. In agreement, BcLOVclust-FUS visibly clustered at 31 °C, and small clusters could be visualized by CluMPS through 34 °C (**Figure 4C, S4C-D**).

In addition to the magnitude of clustering, we found that temperature regulated the dynamics of clustering, such that clustering was less sustained at elevated temperature (T_1/2_ ∼45 min at 31 °C, T_1/2_ ∼75 min around 27 °C with sustained stimulation), consistent with observations from wt BcLOV4 membrane translocation (**Figure 4D-E, Supplementary Movie 2**).

### Multiplexing BcLOVclust and CRY2 for photo-controlled clustering

The availability of multiple optogenetic clustering proteins opens the possibility to cluster multiple targets simultaneously and independently in single cells [19, 20]. To test whether BcLOVclust and Cry2 indeed formed independent clusters, we measured colocalization of BcLOVclust-GFP and mCh-Cry2 under multiple stimulation and expression conditions. Overall, we found no evidence that Cry2 and BcLOVclust interacted within the same clusters. In cells with high expression of BcLOVclust and low levels of Cry2, light induced strong clusters of BcLOVclust but no clusters of Cry2, showing that Cry2 was not being recruited to BcLOVclust (**Figure 5A**). Conversely, in cells with high Cry2 and low BcLOVclust, BcLOVclust was not recruited into Cry2 clusters (**Figure 5B**). In cells with sufficiently high expression of both proteins, light induced clusters in both channels, and the cluster patterns did not overlap (**Figure 5C, Supplementary Movie 3**). To more rigorously assess colocalization, we measured the Pearson correlation coefficient of cytoplasmic fluorescence in the mCh and GFP channels. We compared this metric to ‘positive control’ cells where mCh-Cry2 was additionally fused to a GFP-targeting nanobody [15, 27], and to ‘negative control’ cells where BcLOVclust was coexpressed with mCh alone (**Figure S5**). In cells coexpressing clusters of BcLOVclust-GFP and Cry2-mCh, fluorescence correlation was indistinguishable from the ‘negative control’ cells where mCh was diffuse, further indicating the lack of light-induced association between the two clustering proteins (**Figure 5D**). Thus BcLOVclust and Cry2 can be used to generate spatially distinct light-induced clusters in individual mammalian cells.

**Fig 5.**
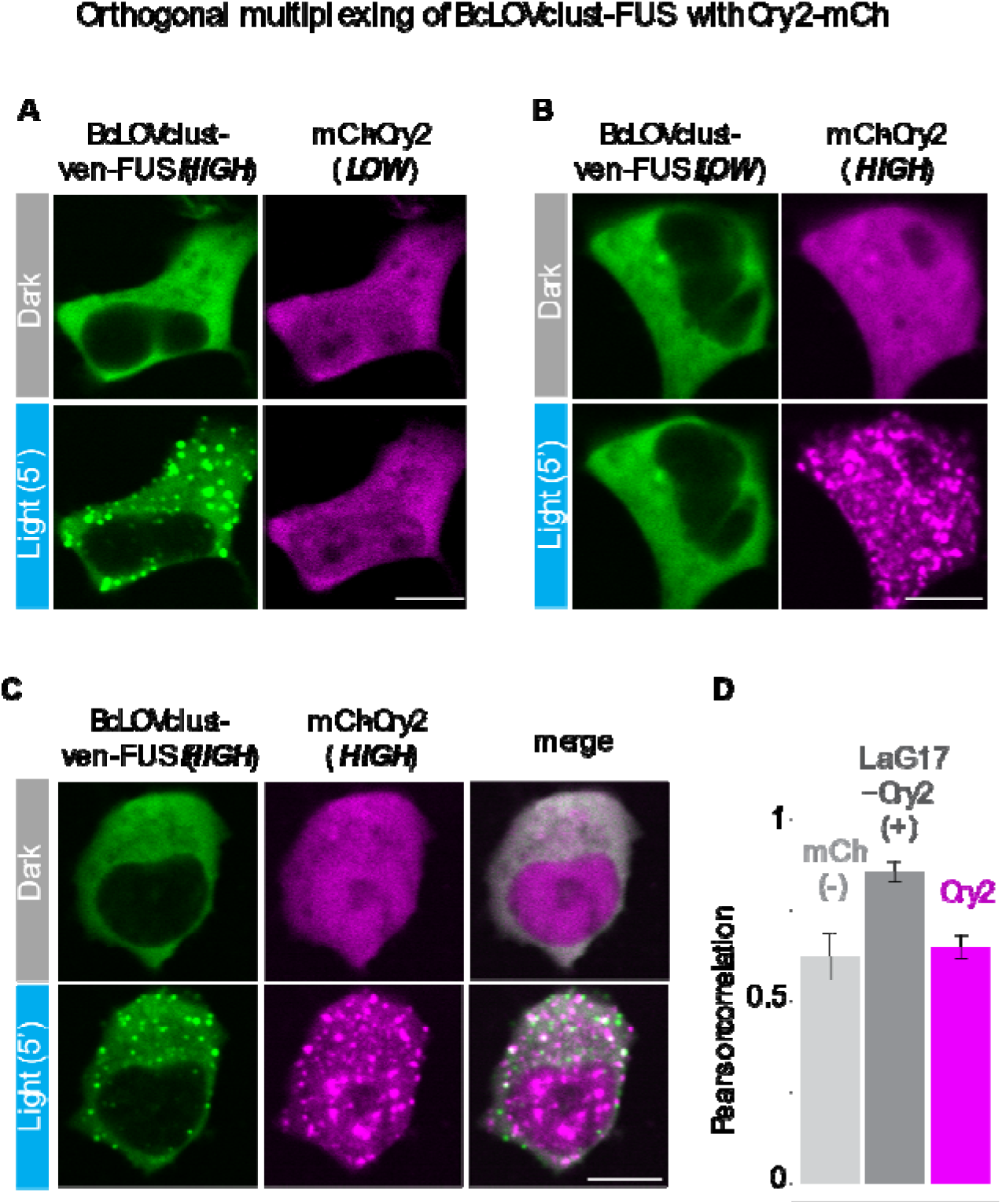
BcLOVclust and Cry2 can be multiplexed to control independent clusters in the same cell. a. In cells with high BcLOVclust-ven-FUS and low Cry2-mCh expression, light induces BcLOVclust clusters that do not recruit Cry2-mCh. b. In cells with high Cry2-mCh and low BcLOVclust-ven-FUS expression, light induces Cry2-mCh clusters that do not recruit BcLOVclust-ven-FUS c. In cells with high expression of both BcLOVclust-ven-FUS and Cry2-mCh, light induces clusters of both constructs that largely do not overlap. For images in (a-c), cells were imaged and stimulated at 25 °C. d. Quantification of the Pearson correlation coefficient of light stimulated BcLOVclust-ven-FUS coexpressed with either 1) mCh, 2) Lag17-mCh-Cry2, or 3) Cry2-mCh. The lowest correlation is observed with coexpression of Cry2-mCh, further suggesting that BcLOVclust and Cry2 clustering do not interact within cells. Cells were stimulated and imaged at 25 °C. More than 100 cells were quantified for each condition. Scale bar, 10 µm.

### BcLOVclust allows control of signaling proteins and stress granules

Finally, we asked whether BcLOVclust could serve as a modular actuator to control intracellular physiology, in a manner similar to Cry2. We first asked whether BcLOVclust could regulate the activity of RhoA, a small GTPase that stimulates cell contractility [13]. In its off state, RhoA resides in the cytoplasm, and its activation is accompanied by translocation to the plasma membrane upon exposure of its C-terminal prenyl group. It was previously found that optogenetic clustering wt RhoA was sufficient to induce translocation and activation to the membrane [13]. Accordingly, clustering of a BcLOVclust-mCh-RhoA fusion induced its localization to the plasma membrane, accompanied by visible cell contraction (**Figure 6A-C, S6, Supplementary Movie 4**). Of note, we also observed that in cells with strong expression, the RhoA fusion appeared partially localized to the membrane in the dark and could be further localized in the dark (**Figure S6A,B**). Expression-dependent basal activity was also observed for Cry2-RhoA, though to a milder degree and only at comparatively higher expression levels (**Figure S6C**).

**Figure 6.**
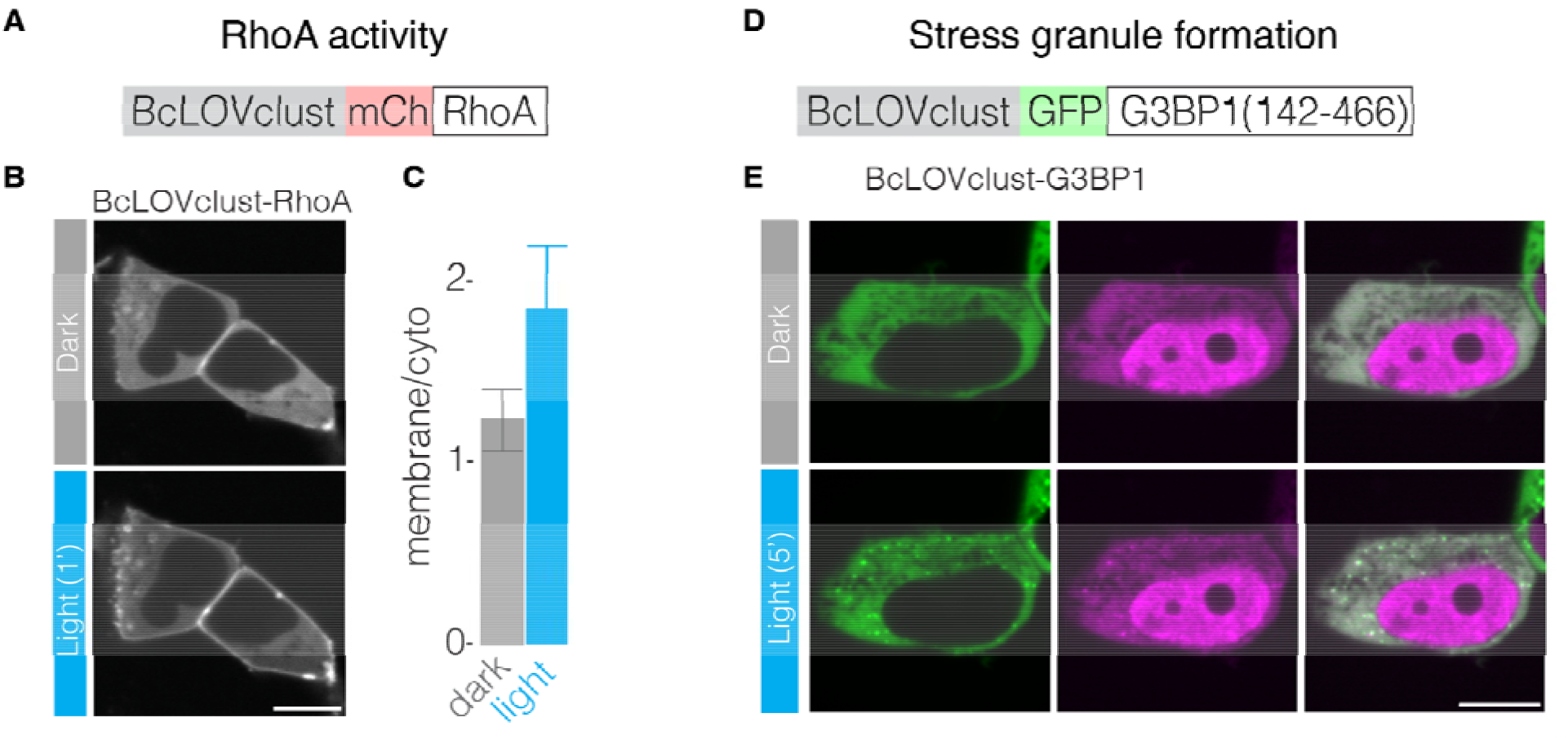
BcLOVclust controls intracellular signaling and stress-associated proteins. a. Domain schematic of BcLOVclust-RhoA. b. Light stimulation of BcLOVclust-RhoA induces its membrane translocation. Cells were stimulated with 3 minutes of 488 nm blue light at 25 C. c. Quantification of translocation in (B) in HEK 293T cells with comparable expression levels. d. Domain schematic of BcLOVclust-G3BP1. E) Light stimulates clustering of G3BP1, the first step of stress granule formation. TIA-1 is a RNA binding protein recruited to stress granules under typical environmental stress as well as light-inducible stress granule formation systems. Scale bar, 10 µm.

As a second example, we asked whether BcLOVclust could regulate the formation of stress granules, as previously shown through the induced oligomerization of a fragment of G3BP1 (amino acids 142-466) [28]. We fused this fragment to BcLOVclust-GFP and, upon expression in HEK 293T cells, observed robust granule formation within 2 minutes of light stimulation. These granules co-localized with the stress granule marker Tia1 (**Figure 6D-E, Supplementary Movie 5**), further indicating that BcLOVclust-G3BP1 was indeed forming functional stress granules with the recruitment of RNA binding proteins and potentially RNA [28]. Together, these demonstrations validate that BcLOVclust is an optogenetic clustering module that could be broadly applied to regulate various phenomena within cells.

## Discussion

By isolating the native clustering properties of the BcLOVclust protein, we have generated BcLOVclust, an optogenetic protein that clusters in the cytoplasm of mammalian cells. BcLOVclust clusters and declusters more rapidly than Cry2, and the extent of clustering can be tuned through several strategies including choice of fluorophore or appending intrinsically disordered regions. Finally we show that BcLOVclust and Cry2 can be multiplexed to form distinct condensates using the same blue light input in single cells, and that BcLOVclust can regulate the activity of intracellular signaling molecules in a modular fashion.

Our work sheds new light on the mechanism of BcLOV4 photoactivation. We previously showed that clustering could tune the membrane-binding potential of wt BcLOV4, though whether membrane binding was required for clustering was not clear [23]. Here our work shows that membrane binding is not required for clustering, and that clustering can be isolated by removing the key basic residues that mediate membrane binding. Nevertheless, our results also suggest limits to this separability, since the 6K1R mutant showed weaker clustering than the 4K mutant, implicating some roles for basic residues in AH1e for clustering. Future work will determine whether different subsets of mutations could allow for stronger retention of clustering while still eliminating membrane binding.

BcLOVclust has distinct functional characteristics compared to Cry2, most notably in its increased speed of clustering and declustering. Beyond allowing better dynamic control over fast processes, these differences could be leveraged to differentially communicate to the two proteins using a single channel of blue light, for example using fast and infrequent pulses to stimulate Cry2 while BcLOVclust remains mostly in its OFF state. Such multiplexed control over distinct condensates could be useful in the study of various systems, for example in nuclear regulation or in the cell stress response, where multiple condensates coexist and together regulate cell behavior [28, 29, 30]. These studies may further benefit from unique strategies to activate either one or both proteins with blue light, for example by specifying sample temperature to promote or suppress BcLOVclust activity.

Despite these features, we also note limitations of BcLOVclust, particularly for use in mammalian cells. BcLOVclust forms light induced clusters only below ∼30 °C. Although compatible with mammalian cell survival, these sub-physiological temperatures may limit the scope and interpretation of experiments. Further studies are required to understand the molecular nature of BcLOV4 thermosensing to engineer variants that will operate at 37 °C. Nevertheless, BcLOVclust should work well in organisms like yeast, flies, and zebrafish, which are cultured at lower temperatures and which can be controlled by wtBcLOV4 [22,26]. BcLOVclust generally forms fewer and smaller clusters compared to Cry2, though several strategies exist to strengthen the degree of clustering, including the two that we demonstrate in this work [9,14,15]. In addition, we observed concentration-dependent basal activation in the dark state of the BcLOVclust-RhoA fusion, although similar basal activation was not observed with BcLOVclust-G3BP1. Thus activation parameters like basal activity and dynamic range must be empirically verified for each application.

## Author contributions

Z.H. and L.J.B. conceived the study. Z.H., W.B., and L.D. performed experiments and analyzed data. L.J.B. supervised the work. Z.H. and L.J.B. wrote the manuscript and made figures.

## Acknowledgements

The authors thank Kevin Gardner for helpful discussions on this work. This work was supported by the National Institutes of Health (R35GM138211 for L.J.B) and the National Science Foundation (Graduate Research Fellowship Program to W.B., CAREER 2145699 to L.J.B.), and the Penn Center for Precision Engineering for Health (CPE4H). Cell sorting was performed on a BD FACSAria Fusion that was obtained through NIH S10 1S10OD026986 and is operated through the Penn Cytomics and Cell Sorting Resource Laboratory.

## Methods

### Molecular cloning

All plasmids were cloned in the pHR backbone with a CMV promoter. Plasmids were assembled using restriction enzyme digestion (MluI and NotI), PCR with Q5 polymerase from NEB (M0493S), followed by HiFi assembly (NEB E2621S) and transformation of chemically competent cells (NEB Turbo, C2984H). BcLOV4 and the FUS-LC region were amplified from previously-descrived constructs [22,23]). Cry2 was amplified from a previously described construct (optoFGFR) [5]. Note that EGFP was used in all “GFP” fusions, and mVenus was used in all “Venus” fusions.

### Cell culture

All studies in the manuscript were carried out in Lenti-X HEK 293T cells. Cells were cultured in DMEM with 10% fetal bovine serum (FBS) and 1% penicillin/streptomycin (P/S). Cells were maintained in standard cell culture incubators at 37 °C with 5% CO_2_ and were moved to other temperatures before each experiment as specified.

### Protein expression

All the BcLOV variants and Cry2 were transiently transfected into HEK 293T cells thatstably expressing iRFP-CAAX or the CluMPS reporter as specified in figures. All the transfections were done using Lipofectamine 3000 (Invitrogen) according to the manufacturer’s protocol. HEK 293T cells were seeded to 40-60% confluence prior to transfection. The CluMPS reporter and iRFP-CAAX cell lines were sorted for stable expression after lentiviral transduction.

### Temperature control

Cells were cultured at 37 °C in normal cell incubators until 2h before an experiment, when they were transferred to incubators or a microscope-associated environmental chambers (Okolab) at the intended experimental temperature. For characterization of BcLOVclust behavior under multiple temperatures **(Figure 4)**, we used a custom, feedback-controlled device device to maintain temperature in wells of a 96-well plate in a manner compatible with imaging through an inverted microscope.

### Confocal imaging

Live-cell imaging was performed using a Nikon Ti2-E spinning disk (Yokagawa CSU-W1) confocal microscope equipped with 488/514/561/640nm laser lines, an sCMOS camera (Photometrics). CO_2_ and ambient temperature were controlled by an environmental chamber (Okolabs). For imaging with the custom device for temperature control, the ambient temperature of the environmental chamber was set to 25 °C. Otherwise, the temperature was set to the temperature specified in figure legends. HEK 293T cells that expressed the construct of interest were imaged with either a 20X or 40X air objective for quantification purposes, and representative images were taken using a 60x oil objective, at variable temperatures and 5% CO_2_. BcLOV4 variants were stimulated using the 488 nm laser.

All imaging experiments were performed using the same field of view along the time course except for **Figure 2B and 3E**, which measured BcLOVclust declustering kinetics. We observed that the 514 nm laser used to image mVenus could weakly stimulate BcLOVclust. To best measure declustering kinetics, we thus avoided continuous imaging of EGFP and mVenus in order to avoid unintended stimulation. Instead, we imaged different fields for each time point of dark adaptation after initial stimulation of all fields. In other words, each field was imaged only once after the stimulation phase to record the remaining clusters.

### Image analysis

#### Membrane recruitment

The membrane association of BcLOV4 and its variants in **Figure 1E** was calculated using normalized plasma membrane (PM)/cytoplasmic mean fluorescence ratio. Cells were segmented using an iRFP-CAAX (Kras) marker (**Figure S7**), which localizes to the plasma membrane, using the MorpholibJ module in ImageJ. These segmented images were then imported into a CellProfiler script, in which the cells with positive GFP signal and proper area were selected. The PM is defined by a 3 pixel ring inside the edge of the contour of each cell. The PM and whole cell fluorescence were then quantified.

#### Clustering

Fluorescent clusters were quantified using a custom MATLAB script that identified pixels having steep contrast over the surrounding fluorescence signal. Briefly, we first segmented the cells by manually selecting cells based on their fluorescence (not including nucleus). Then, we passed these segmentations to different time points and fluorescence channels and modified them as necessary. Next, we identified clusters within each cell in a semi-automated manner through an iterative adjustment of two parameters that maximized true cluster identification while suppressing noise. Finally, we exported the pixel-level data and classification and analyzed cluster intensity, area, number per cell in R. Matlab and R code are available upon request (**Figure S8**).

#### Colocalization

The colocalization between 2 different fluorescent signals in this manuscript is quantified using pearson correlation. Since BcLOVclust is mainly localized to the cytosol, we only used cytosolic fluorescence for the calculations (**Figure 5D**).

### Curve fitting, normalization and plotting

To generate fits of photoreceptor clustering over time (**Figure 2A,B**, **Figure 3 D,E**, and **Figure S1**), we used exponential decay for BcLOV variants (**Eq.1**) and sigmoidal fits using the Hill equation (**Eq.2**) for Cry2 data. These distinct fitting functions best minimized error for the two photoreceptors.

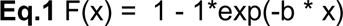

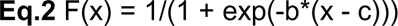

All the fitting curves were rescaled between 0 and 1 according to the minimum and maximum values of the data points. **Figure 1E** is normalized to the dark state (1st time point) for visualization.

## Supplementary Figures

**Fig S1.**
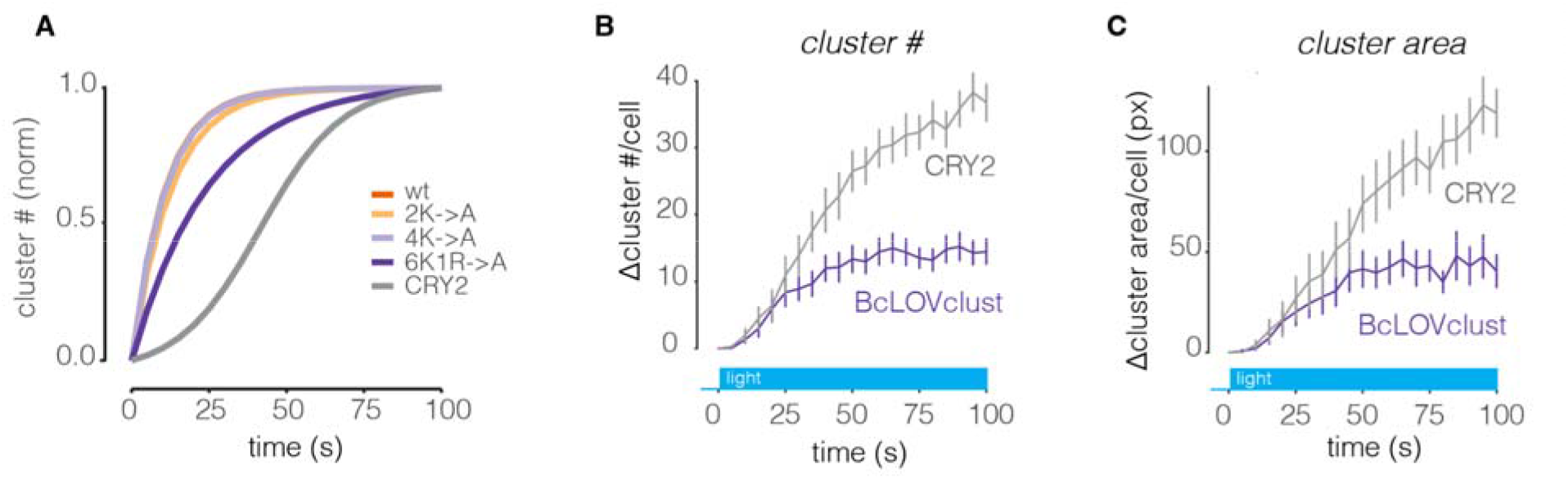
Clustering kinetics of all BcLOV variants tested are faster than Cry2. a. To compare kinetics of stimulation, all data in **Figures 1F** and **2A** were fit to either exponential or Hill equations, rescaled between 0 and 1, and overlaid. b. Comparison of absolute cluster number from **Figure 2A**. c. Comparison of absolute cluster area from **Figure 2A**

**Fig S2.**
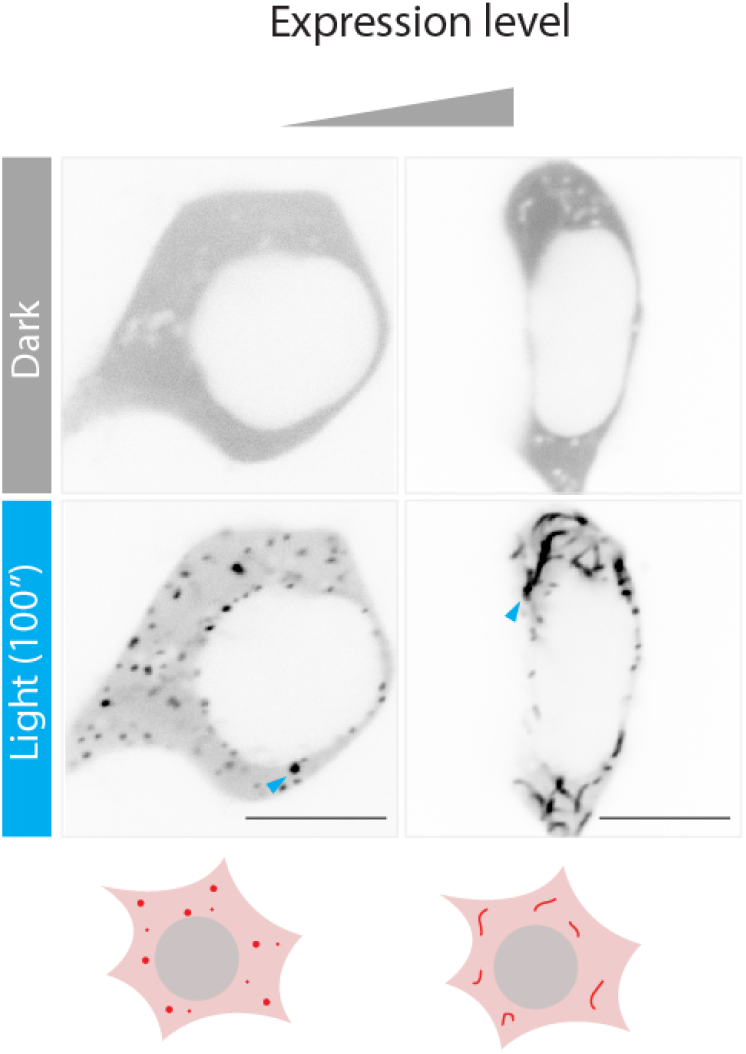
N-terminal fusion of FUS-LC to BcLOVclust forms light-induced fibers in a concentration-dependent manner. Representative images of HEK 293T cells expressing FUS-BcLOVclust-mCh, imaged at 25 °C. Image brightness was adjusted between the “low” and “high” conditions for clearer comparison of clusters and fibers. Cells were stimulated every 10 seconds. Scale bar, 10 µm.

**Fig S3.**
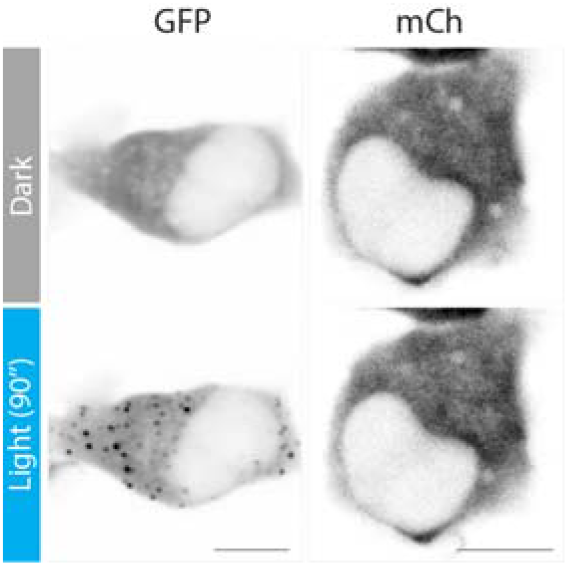
Choice of fluorescent protein alters the magnitude BcLOV clustering. BcLOVclust was fused to either mCherry or GFP at the C-terminus. Images show stronger clustering with GFP than with mCherry. Cells were stimulated with 488 nm light every 15 s at 25 °C. Scale bar, 10 µm.

**Fig S4.**
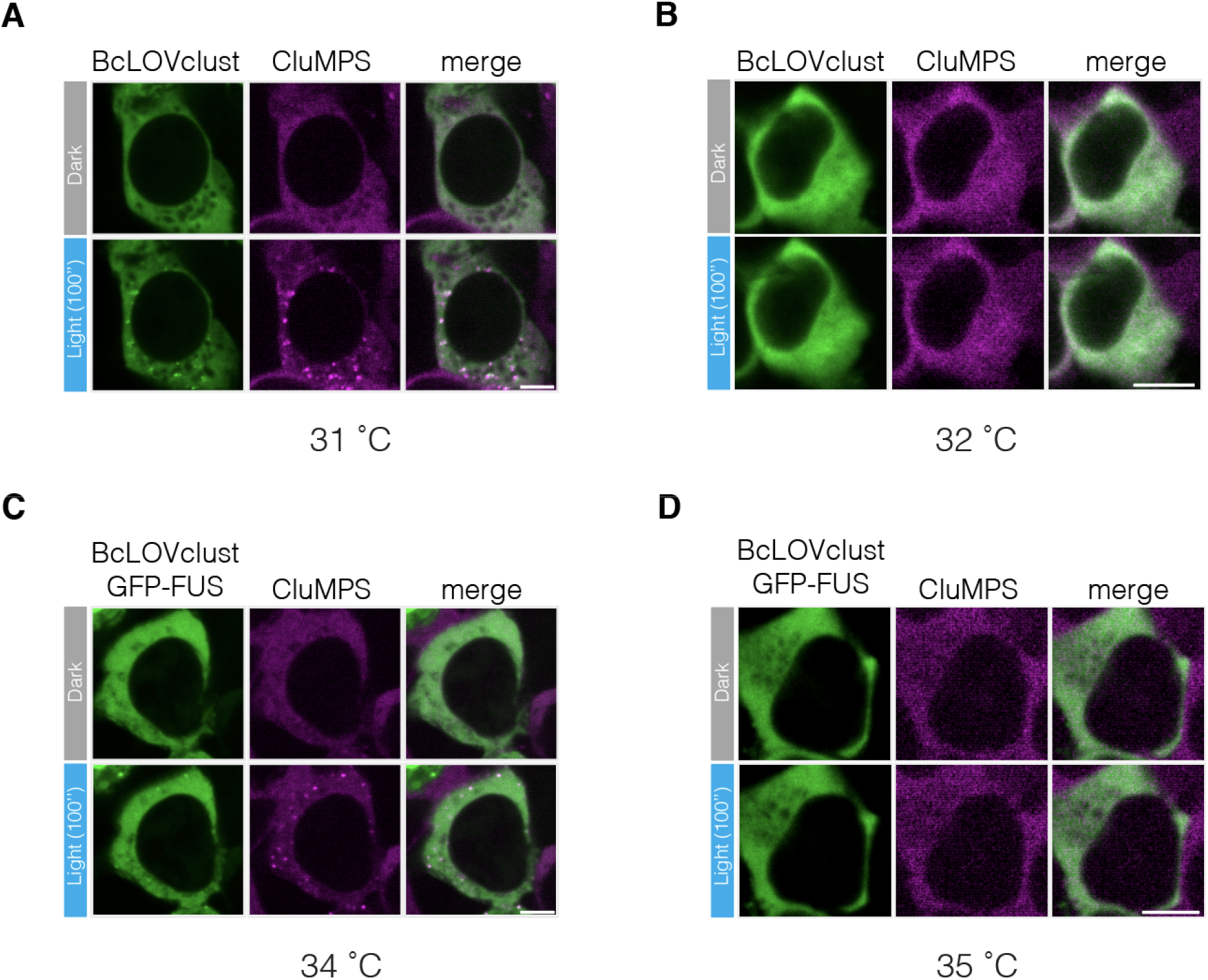
FUS-LC permits clustering at higher temperatures. a. Representative image of small BcLOVclust-GFP clusters forming at 31 °C in HEK 293T cells. Clusters were amplified by co-expression of a CluMPS reporter. b. These clusters were no longer visible at 32 °C. c. Representative image of small BcLOVclust-GFP-FUS clusters forming at 34 °C in HEK 293T cells. Clusters were amplified by co-expression of a CluMPS reporter. d. These clusters were no longer visible at 35 °C. Scale bar, 10 µm.

**Fig S5.**
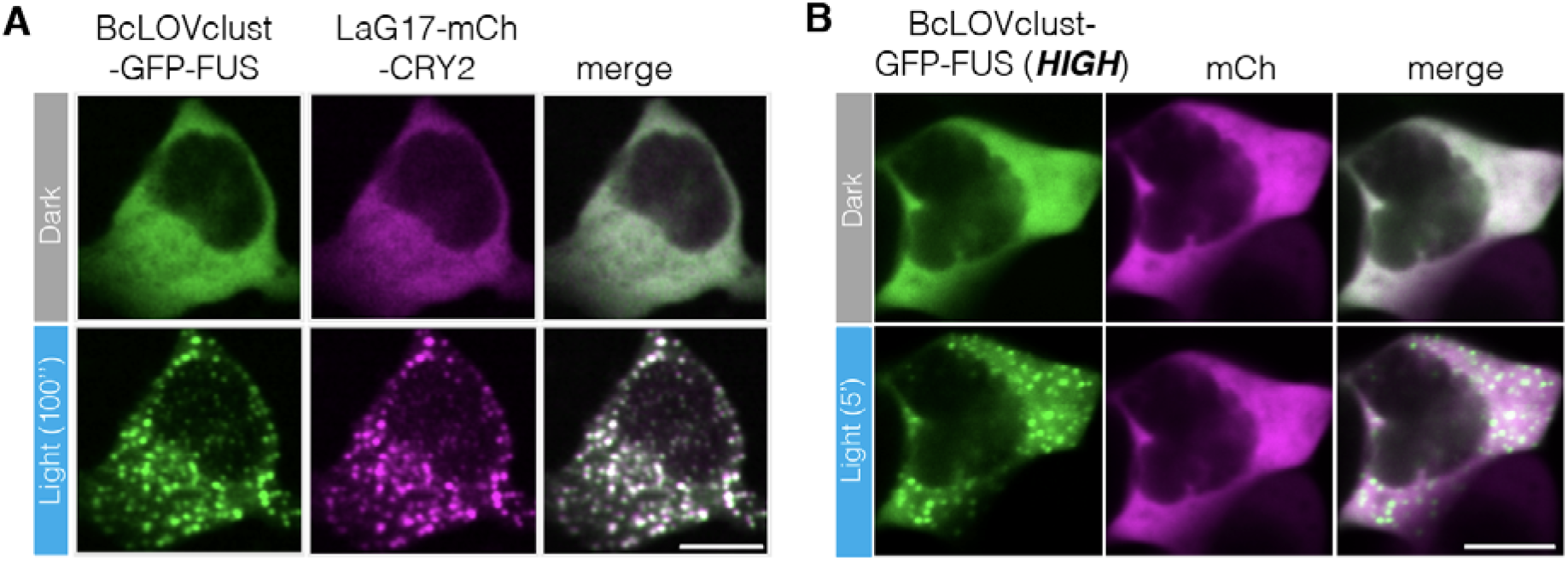
Positive and negative controls for measuring colocalization of BcLOVclust and Cry2 clusters. a. Representative images of the positive control for colocalization. BcLOVclust-GFP-FUS was cotransfected with mCh-Cry2 fused to a nanobody for GFP (LaG17). Blue light induced clustering of both BcLOVclust and Cry2, and clusters colocalized with each other due to affinity between GFP and Lag17. b. Representative images of the negative control for colocalization. BcLOVclust-GFP-FUS was cotransfected with mCherry. Blue light induced clustering of only BcLOVclust. Experiment was performed at 25 °C. Scale bar, 10 µm.

**Figure S6.**
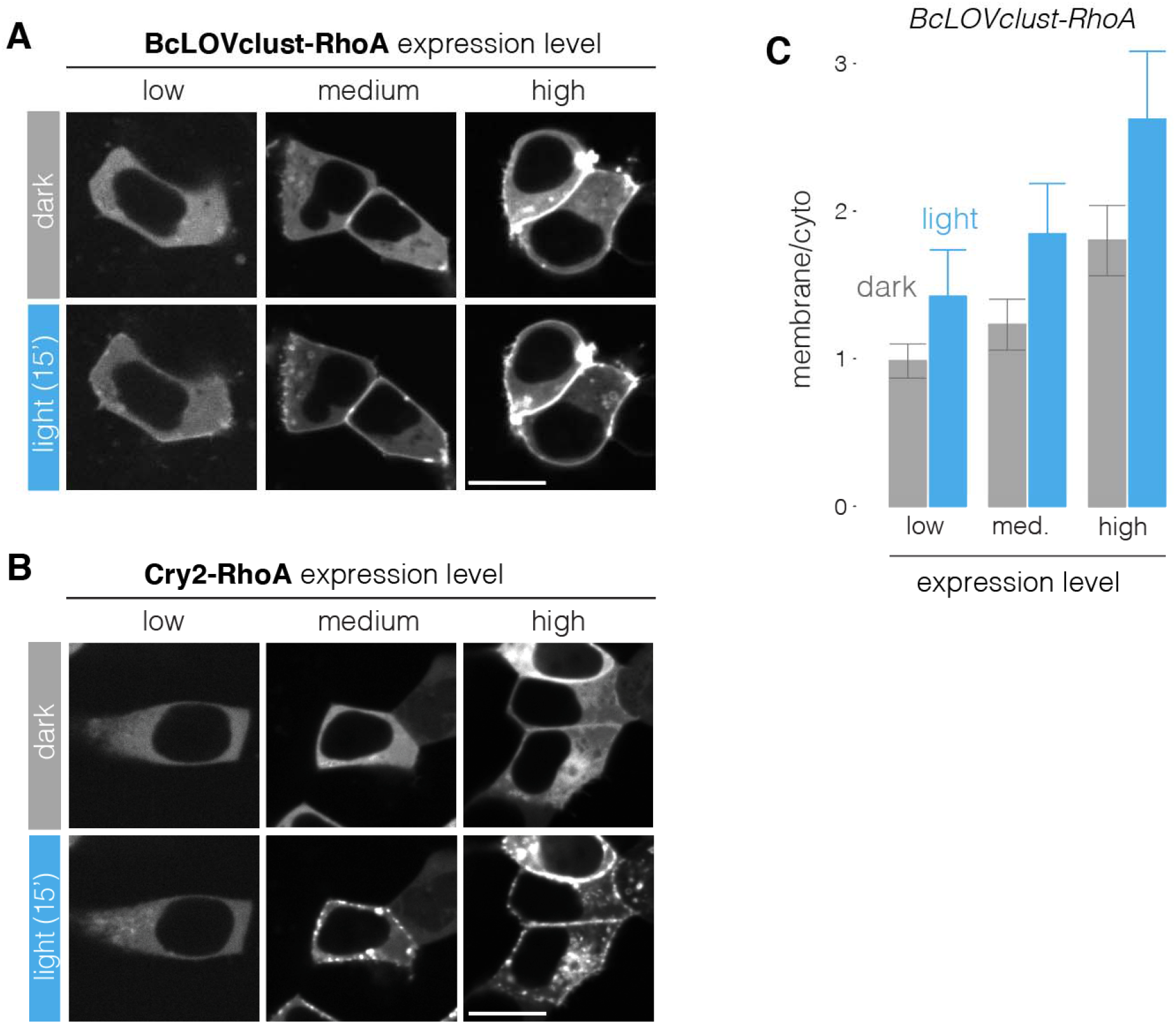
Dark-state and lit-state activation of BcLOVclust-RhoA depends on its concentration. a. Representative images of BcLOVclust-RhoA activation in HEK 293T cells, performed at 25 °C. While light induces clustering and translocation of RhoA, dark-state association with the membrane can be observed at elevated expression levels. b. Quantitation of experiment shown in (a). Bars represent the ratio of mean membrane to cytoplasmic pixel intensity. Each bar represents the mean ratio from 10 cells. Error bars represent standard deviation. c. Relative to BcLOVclust-RhoA, Cry2-RhoA shows milder expression-dependent basal activation, and only at the highest expression levels. Scale bars, 15 µm.

**Fig S7.**
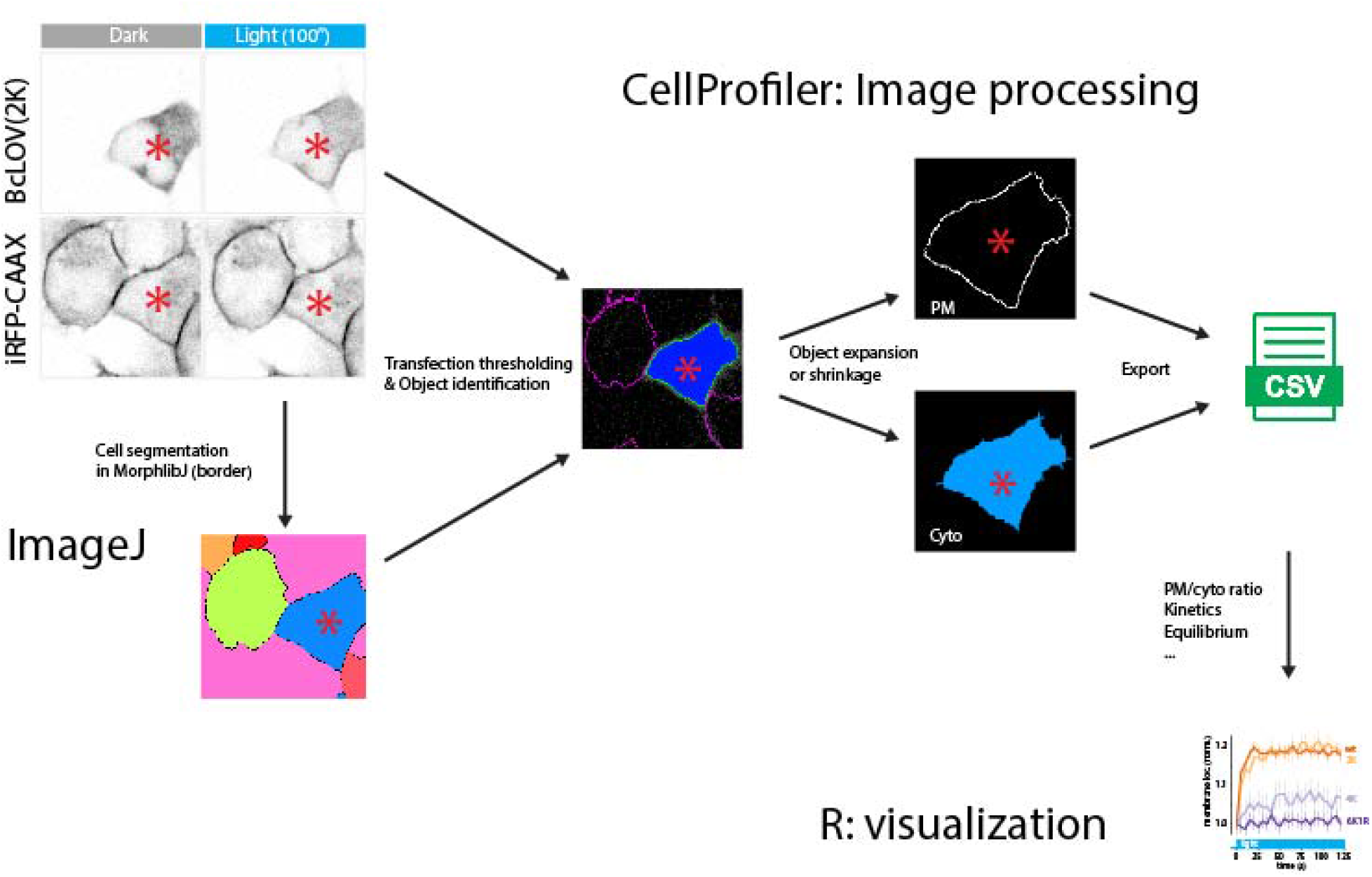
Image analysis pipeline for quantifying PM recruitment of proteins. Target constructs (BcLOV(2K->A) shown as an example) were transfected in HEK 293T cell lines that expressed a marker for the plasma membrane (here, iRFP-CAAX). The iRFP channel was used to segment cells by MorphoLibJ in ImageJ. Of these segmented cells, cells that expressed BcLOV-GFP analyzed in CellProfiler. These objects were either expanded or shrunken to identify pixels that correspond to the plasma membrane or cell cytoplasm, respectively. Pixel values were quantified and visualized plotted in R.

**Fig S8.**
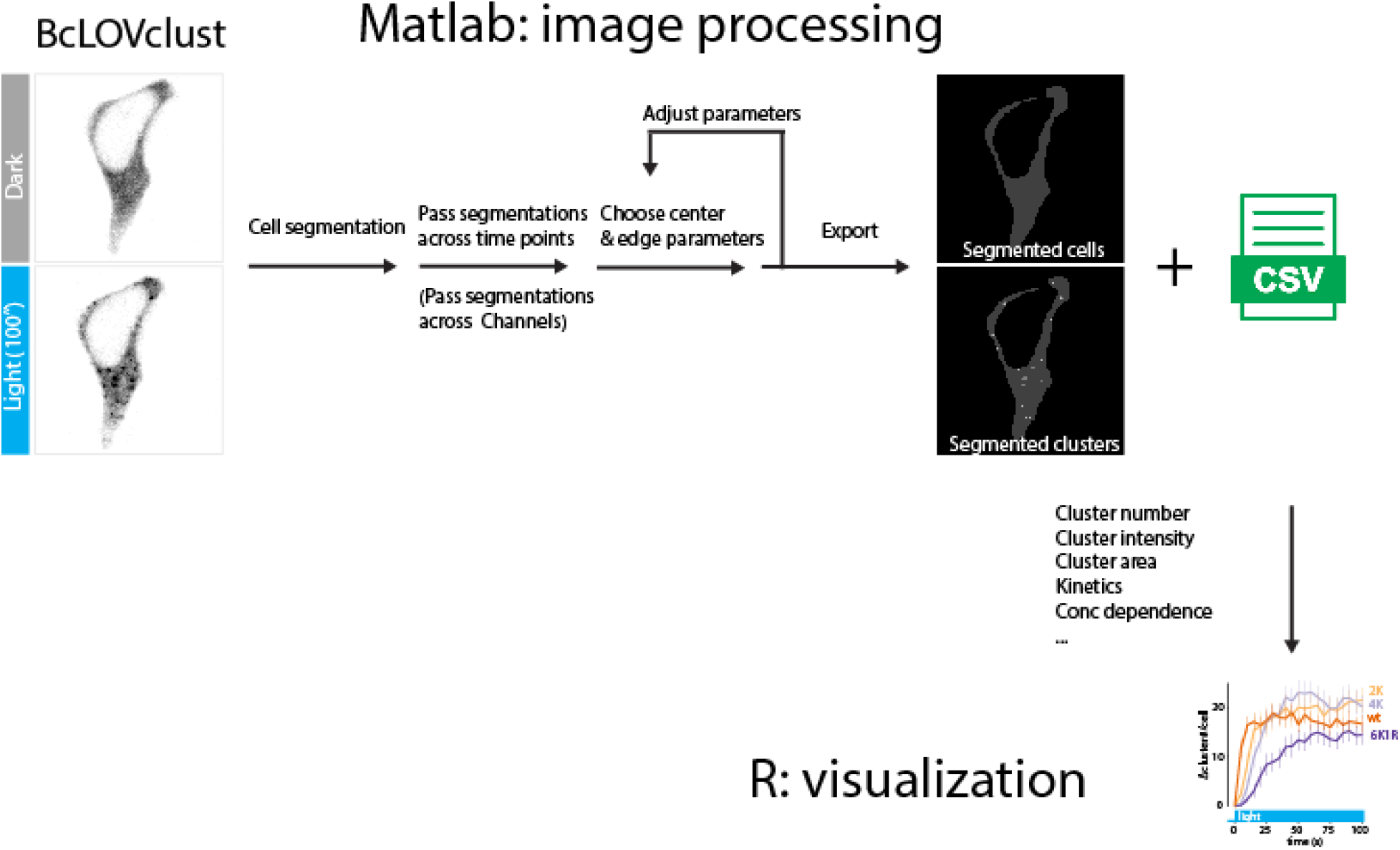
Image analysis pipeline for quantification of protein clusters. Target constructs (BcLOVclust as an example) were transfected into HEK 293T cells and imaged using a confocal microscope. The images were loaded into a MATLAB pipeline where cells are segmented manually. The cell segmentations are then passed to different time points and channels, and clusters are identified in a semi-automated fashion by specifying two parameters (center and edge values). Cluster number and area are derived from the counts and intensities of the pixels within these identified clusters.

## Supplementary Movie Captions

**Supplementary Movie 1. Removal of positive charges from AH1e progressively weakens membrane binding but retains homo-oligomerization.** HEK 293T cells expressing the indicated BcLOV4 variant fused to GFP were imaged and stimulated with 488 nm laser light at 29 °C. Time is mm:ss, scale bar is 10 µm.

**Supplementary Movie 2. BcLOVclust clusters are transient at higher temperatures.** HEK 293T cells co-expressing BcLOVclust-GFP and a GFP-targeting CluMPS reporter were imaged and stimulated with 488 nm laser light at the indicated temperatures. GFP channel is shown. Time is hh:mm:ss, scale bar is 10 µm.

**Supplementary Movie 3 Multiplexing BcLOVclust and Cry2 in individual cells.** HEK 293T cells co-expressing BcLOVclust-Venus and a mCh-Cry2 were imaged and stimulated with 488 nm laser light at 25 °C. Time is mm:ss, scale bar is 10 µm.

**Supplementary Movie 4 Light-induced stimulation of BcLOVclust-RhoA.** HEK 293T cells expressing BcLOVclust-GFP-RhoA were imaged and stimulated with 488 nm laser light at 25 °C. Arrow indicates region of visible light-induced contractility, indicative of RhoA activity. Time is mm:ss, scale bar is 10 µm.

**Supplementary Movie 5 Light-induced stimulation of BcLOVclust-G3BP1.** HEK 293T cells co-expressing BcLOVclust-GFP-G3BP1 and the stress granule-associated protein mCh-TIA1 were imaged and stimulated with 488 nm laser light at 25 °C. Light induces clusters of BcLOVclust that colocalize with mCh-TIA1. Time is mm:ss, scale bar is 10 µm.

